# Tier-specific location of Lewy body pathology and related neuromelanin levels drive dopaminergic cell vulnerability in pigmented non-human primates

**DOI:** 10.64898/2026.03.30.715197

**Authors:** Julia Chocarro, Alberto J. Rico, Goiaz Ariznabarreta, Elena Lorenzo-Ramos, Mario M. Ilarduya, Cristina Canales, Adriana Leon-Villares, Javier Blesa, José A. Obeso, José L. Lanciego

**Affiliations:** CNS Gene Therapy Program, Center for Applied Medical Research (Cima), University of Navarra, Pamplona, Spain; Centro de Investigación Biomédica en Red de Enfermedades Neurodegenerativas (Ciberned-ISCIII), Madrid, Spain; Centro Integral de Neurociencias Abarca Campal (HM-CINAC), Hospital Universitario HM Puerta del Sur, HM Hospitales, Madrid, Spain; Instituto de Investigación Sanitaria HM Hospitales, Madrid, Spain; Facultad HM de Ciencias de la Salud, Universidad Camilo José Cela, Madrid, Spain

**Author notes:** Corresponding author: *<u>José L. Lanciego, MD, PhD</u>, CNS Gene Therapy Program, Center for Applied Medical Research (CIMA), University of Navarra., Pio XII Avenue 55, 31008 Pamplona (Navarra), Spain, E-Mail:.

**Keywords:** alpha-synuclein, tyrosinase, substantia nigra, adeno-associated viral vectors, neurodegeneration

## Abstract

Although a differential vulnerability of dopaminergic neurons to degeneration based on their specific location within the dorsal and ventral tiers of the substantia nigra pars compacta (SNcD and SNcV, respectively) has long been postulated, the underlying mechanisms sustaining these tier-specific differences remain poorly understood. Here, upon inducing a viral-mediated enhancement of neuromelanin (NMel) accumulation within dopaminergic neurons in non-human primates, the distribution of Lewy body-like inclusions (LBs) was analyzed within identified SNcD and SNcV neurons, together with their intracellular NMel levels. Results showed that the vast majority of intracytoplasmic inclusions were found in SNcV neurons, and indeed correlated to higher pigmentation levels. By contrast, only very few LBs were found in calbindin-positive neurons of the SNcD, which in parallel exhibited very low levels of NMel accumulation. These results postulate an additive effect made of a tier-specific location of LB burden together with high pigmentation levels as synergistic drivers sustaining the preferential vulnerability of SNcV dopaminergic neurons. Moreover, the evidence obtained here supported that NMel accumulation beyond a given threshold triggers the aggregation of endogenous α-Syn in the form of LBs; therefore, approaches intended to reduce pigmentation levels in SNcV neurons would likely induce a neuroprotective effect by preventing the subsequent aggregation of α-Syn.

## INTRODUCTION

Lewy bodies (LBs) are intracytoplasmic inclusions that typically characterize brain disorders such as Parkinson’s disease (PD) and dementia with Lewy bodies (DLB), and are considered the characteristic histopathological signature of synucleinopathies^1^. The absence of animal models of PD that accurately reproduce LB burden in the SNc represents a significant barrier to the development of novel therapeutic candidates for PD, particularly those focused on disease-modifying interventions^2^. The recent introduction of rodent and non-human primate (NHP) models of PD based on the delivery of viral vectors coding for the human tyrosinase gene (hTyr) has led to pigmentation levels similar to those observed in elderly humans and can be considered reliable choices that overcome this limitation^3–5^. Animal models based on enhanced NMel accumulation mimic the known neuropathological hallmarks of PD with unprecedented accuracy, including ongoing pigmentation, presence of LBs made of endogenous aggregates of α-Syn, progressive dopaminergic cell death, and a microglial-driven pro-inflammatory scenario^3–6^.

Within the context of PD, a tier-specific pattern of dopaminergic cell vulnerability has been described, in which calbindin-positive neurons of the dorsal tier of the SNc (SNcD) are more resilient to degeneration than calbindin-negative neurons of the ventral tier (SNcV) ^7–16^. These findings paved the way for appointing calbindin (CB) as a potential neuroprotective agent by buffering calcium overload and thereby protecting dopaminergic neurons against PD-related neurotoxicants^9–10,17–20^.

In keeping with the dichotomy of tier-specific dopaminergic cell vulnerability, a non-random distribution pattern of LBs has been found here in pigmented NHPs. Intracytoplasmic LBs have been almost exclusively found in heavily neuromelanized, calbindin-negative neurons of the SNcV, whereas SNcD neurons only accounted for a minimal fraction of LBs. Accordingly, a synergistic scenario driving dopaminergic cell vulnerability that comprises pigmentation levels, the lack of calbindin, and the tier-specific location of LB pathology, is postulated here.

## RESULTS

Intraparenchymal deliveries of AAV1-*hTyr* in the left SNc resulted in NMel levels high enough to allow direct macroscopic visualization of the pigmented SNc in all experimental subjects, with a trend for a time-dependent slight reduction of NMel levels in animals with 8 months in-life, better observed in female subjects (Figure 1). At the microscopic level, NMel is visualized as a brown-to-black granular pigment occupying most of the cytoplasmic space (Figure 2). Aiforia®-based quantification of pigmented dopaminergic neurons estimated that the viral-mediated enhanced expression of human tyrosinase resulted in pigmentation of roughly up to 40% of the total number of TH+ neurons^4^.

**Figure 1:**
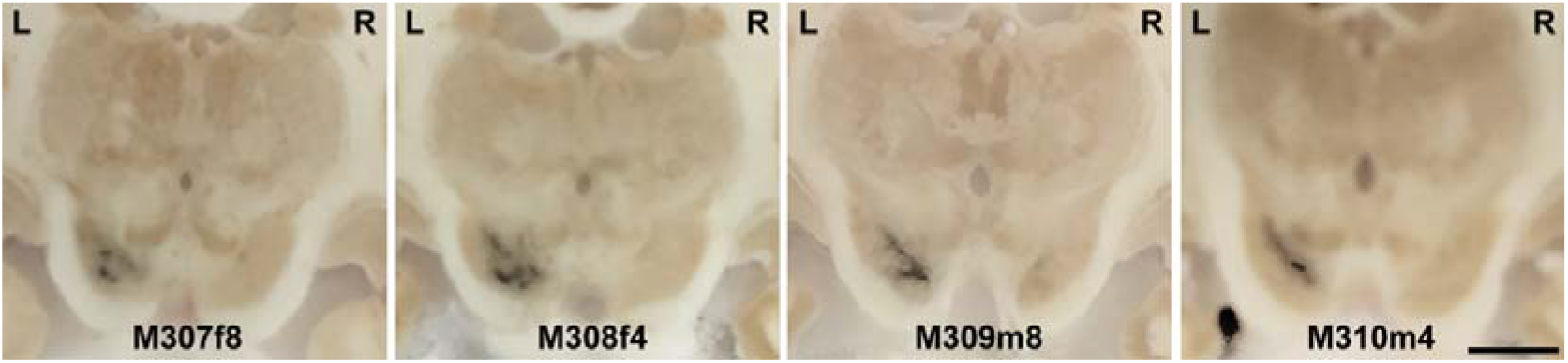
Macroscopic pigmentation of the SNc. Following the delivery of AAV1-*hTyr* into the left SNc, the obtained accumulation of NMel was high enough to allow the direct visualization of the pigmented SNc when the brains were placed in the sliding microtome for sectioning. Scale bar = 3,000 μm

**Figure 2:**
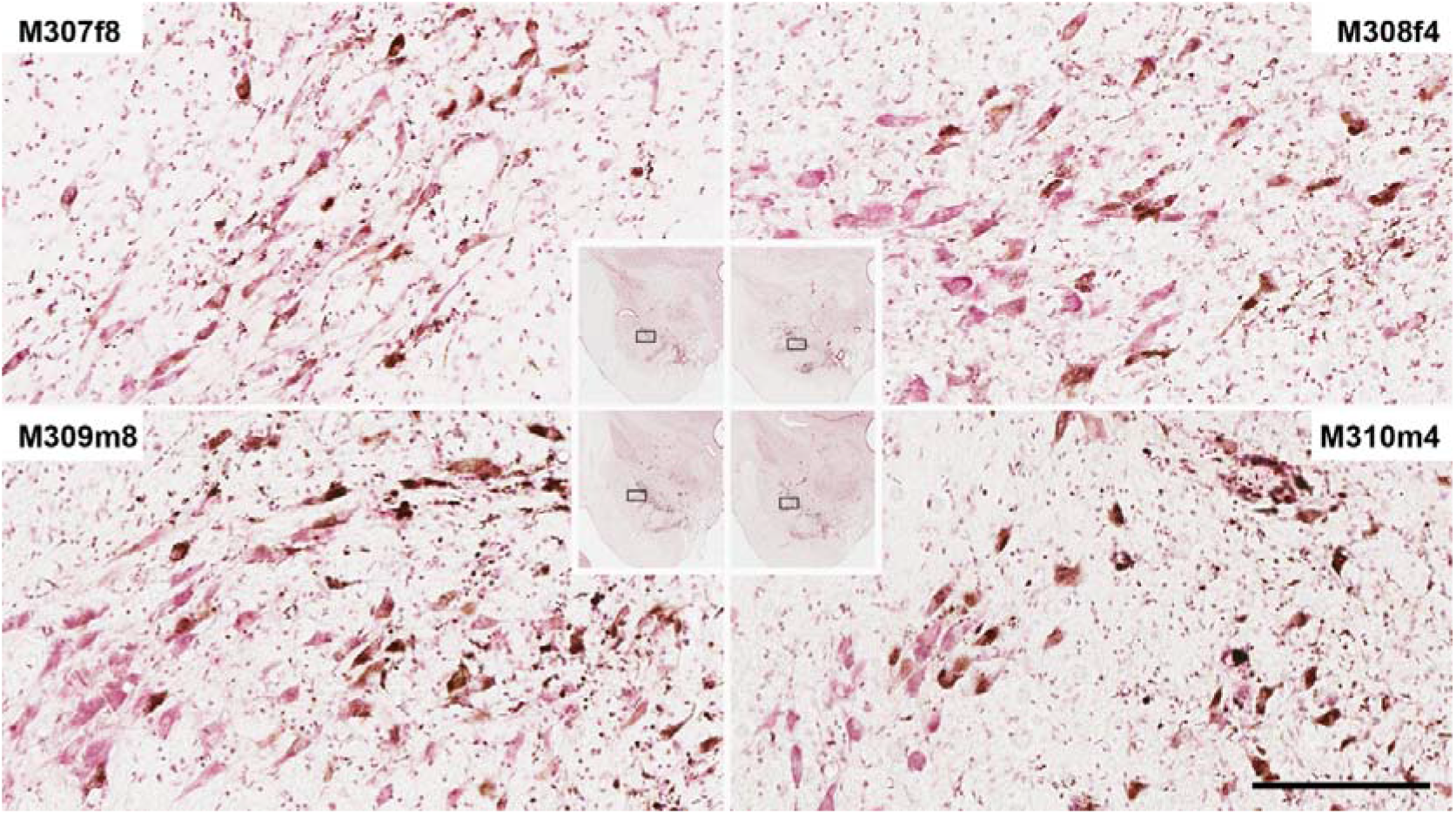
Microscopic pigmentation of SNc neurons. Illustrative photomicrographs taken from sections counterstained with neutral red to disclose between pigmented and non-pigmented neurons in all experimental subjects. Scale bar is 3,000 μm in low-power photomicrographs and 150 μm in high-magnification insets.

### Pigmentation patterns of dopaminergic neurons are tier-specific

In keeping with available evidence^16,21^, pigmented neurons were assigned to either the SNcD or the SNcV based on their differential expression of CB or Aldh1a1 proteins, respectively. The conducted immunoperoxidase stains revealed that a vast majority of pigmented neurons were located in SNc territories lacking calbindin expression (Figure 3). A detailed quantification of both neurochemical endophenotypes showed that in all animals (males and females, and with either four or eight months in-life), pigmented ALdh1a1+ SNcV neurons are the predominant phenotype, accounting for up to 85% of the total number of pigmented neurons on average, whereas the numbers of CB+ neurons (SNcD neurons) only reached up to 15% of pigmented neurons (Figure 3). Furthermore, important phenotype-specific differences were found regarding intracellular NMel levels, since Aldh1a1+ neurons in the SNcV exhibited more abundant intracytoplasmic NMel accumulation than pigmented CB+ neurons located in the SNcD across all experimental subjects (Figures 3A-3C). Finally, it is also worth noting that lower intracellular pigmentation levels for CB+ neurons were found in male specimens compared to those observed in female animals (*p* = 0.001). In summary, in terms of neuromelanized dopaminergic neurons, SNcV neurons are those most often pigmented and indeed exhibiting higher NMel intracytoplasmic accumulation than CB+ SNcD neurons.

**Figure 3:**
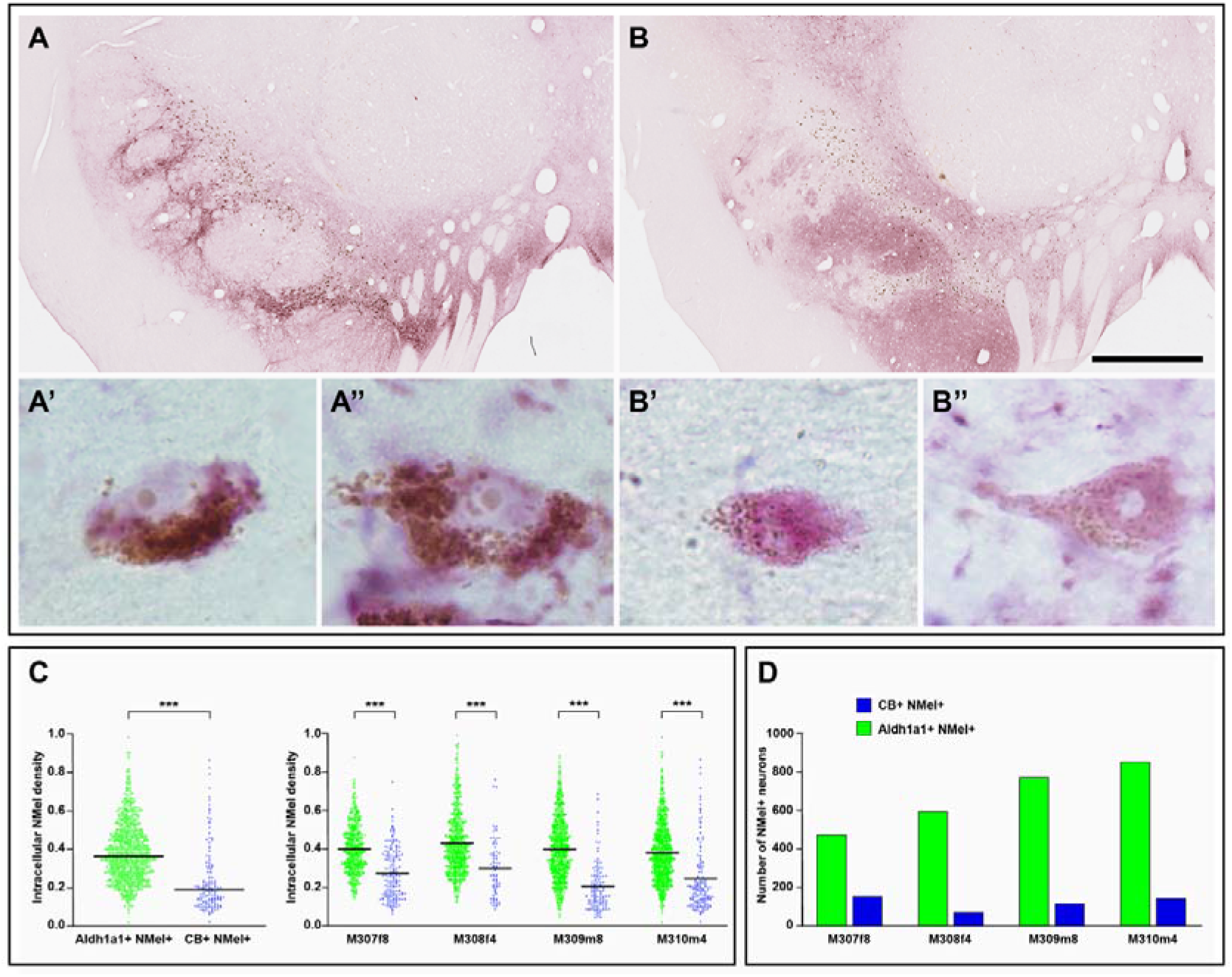
Tier-specific patterns of pigmentation. Immunoperoxidase stains for Aldh1a1 (panels A-A’’) and for CB (panels B-B’’) were conducted to disclose ventral and dorsal tiers of the SNc (SNcV and SNcD, respectively). Pigmented neurons exhibited a preferential pattern of distribution within the SNcV, most of them located in CB-negative territories (panels A and B). On average, 85% of pigmented neurons are located in Aldh1a1+ territories, whereas only up to 15% of neuromelanized neurons are found in the SNcD (panel D). In addition to such a tier-specific pattern of distribution, the intracellular pigmentation levels within SNcV neurons are by far much higher than those found in SNcD neurons (panels A’-A’’, B’-B’’, C). Quantification of intracellular NMel levels was conducted at the single-cell level using Fiji ImageJ software, and comparisons were made using a linear mixed-effects model including dorsal/ventral tier as fixed effects and subject as a random effect accounting for clustering of neurons within subjects, which resulted in NMel+ Aldh1a1+ neurons showing significantly higher OD values than NMel+ CB+ neurons (*p* < 0.001). Two-tailed unpaired *t*-tests performed within individual specimens maintained statistical significance (*p* < 0.001). Scale bar is 1.500 μm in panels A and B.

### Tier-specific location of LB-like inclusions

Intracytoplasmic inclusions positive for traditional LB markers such as phosphorylated α-Syn (pSer129) and P62 were often found within pigmented, TH+ neurons in all experimental subjects. Observed inclusions exhibited ring-shaped morphological characteristics very similar to those reported for LBs in post-mortem human material, such as a peripheral, darker halo and a paler core. They were often found near the transition between the nucleus and the cytoplasm (Figure 4). As expected, LBs were never found in non-pigmented TH+ neurons.

**Figure 4:**
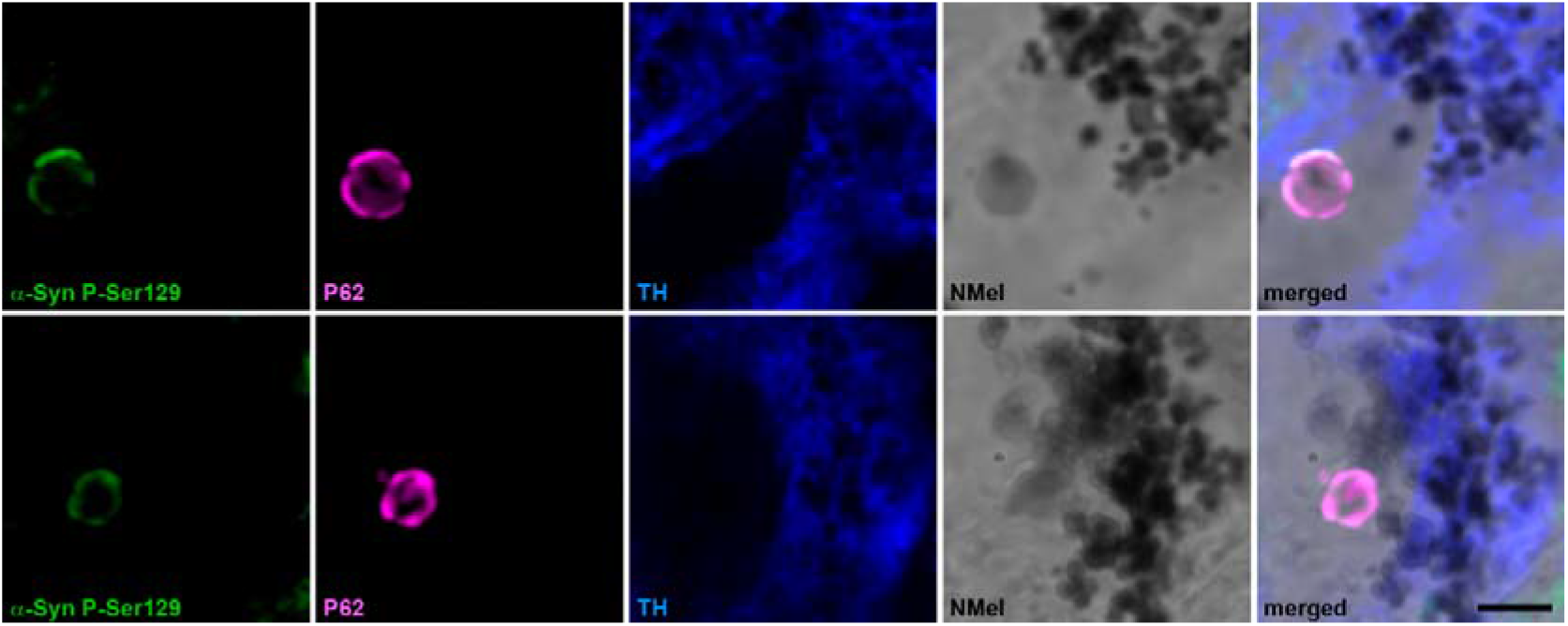
Intracytoplasmic inclusions. Illustrative examples of LBs within neuromelanized dopaminergic neurons. Observed inclusions exhibited ring-shaped morphological characteristics very similar to those reported for LBs in post-mortem human material, such as a peripheral, darker halo and a paler core. They were often found near the transition between the nucleus and the cytoplasm. Inclusions were immunoreactive for traditional markers of LB burden, such as phosphorylated α-Syn (P-Ser129; green channel) and P62 (magenta channel). Scale bar is 5 μm.

From the total number of neuromelanized neurons, approximately two-thirds of pigmented neurons lacked LB inclusions (mean value of 64.26%), whilst intracytoplasmic aggregates have been observed in the remaining one-third (36.11% on average) (Table1). Within pigmented neurons showing LB burden, a precise tier-specific location has been found (Figure 5) with ventral tier neurons (Aldh1a1+) showing the highest percentage of inclusions (95.65%) compared to a marginal 4.33% of pigmented CB+ neurons with LB pathology (Figure 6A).

**Table 1:** Tier-specific location of LB pathology within pigmented neuronal endophenotypes.

| Cell type | M307f8 | M308f4 | M309m8 | M310m4 | Mean value |
| --- | --- | --- | --- | --- | --- |
| NMel+ P62- | 70.85% | 63.95% | 60.10% | 62.14% | 64.26% |
| NMel+ P62+ Aldh1a1+ | 29.14% | 36.55% | 37.94% | 34.86% | 34.52% |
| NMel+ P62+ CB+ | 0.8% | 1.52% | 1.95% | 2.11% | 1.59% |

**Figure 5:**
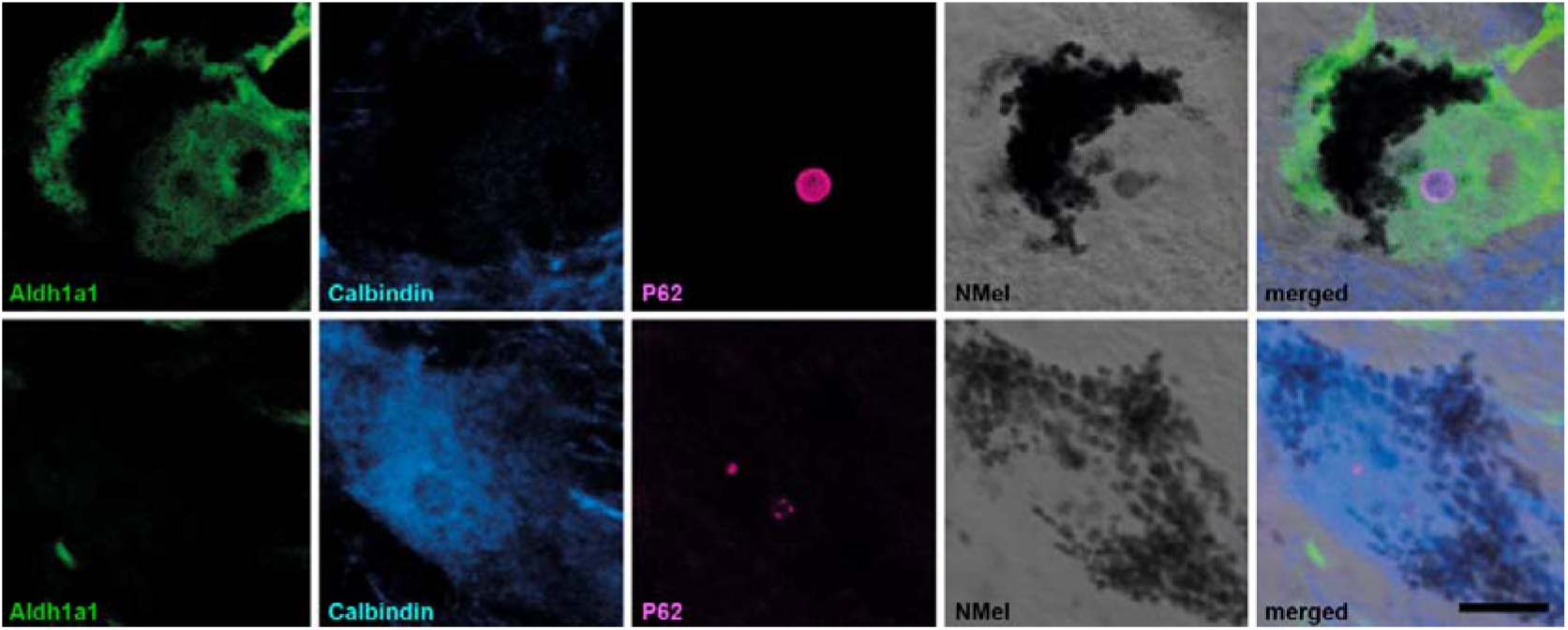
Tier-specific distribution of intracytoplasmic inclusions. The vast majority of LBs were consistently found within heavily pigmented dopaminergic neurons located in the ventral tier of the SNc (Aldh1a1+ neurons; green channel). By contrast, LBs were rarely observed within calbindin-positive neurons of the dorsal tier (blue channel), likely resulting from the weaker pigmentation that typically characterizes this neurochemical subtype of dopaminergic neurons. Scale bar is 10 μm.

**Figure 6:**
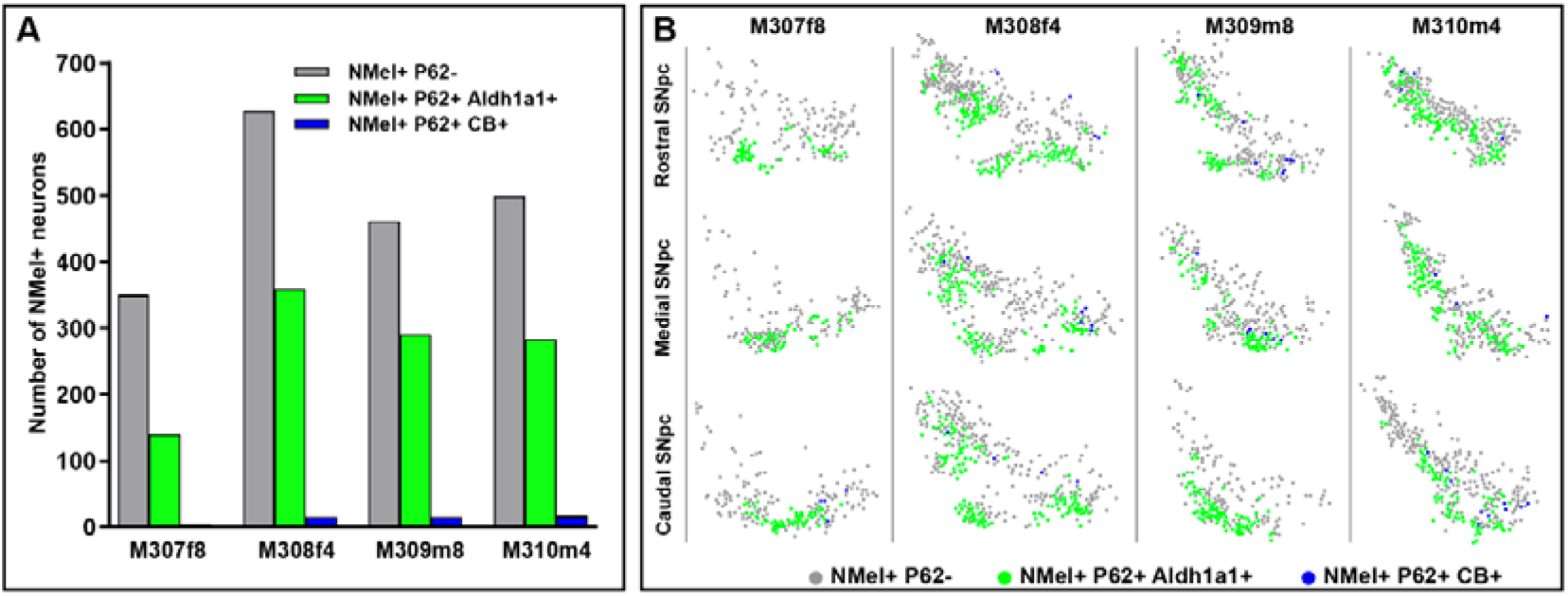
Tier-specific patterns of LBs location. (A) Histograms showing the number of pigmented dopaminergic neurons across all experimental subjects, categorized by their tier-specific location and the presence or absence of LBs. Roughly about one-third of the pigmented neurons exhibited intracytoplasmic inclusions (P62+). (B) Location of LBs in each experimental animals, as seen in three coronal levels covering the whole rostrocaudal extent of the SNc (rostral, medial, and caudal). A vast majority of LBs inclusions were found in pigmented Aldh1a1+ neurons (95.65%), whereas a minimal fraction of LBs were observed in CB+ neurons (4.33% on average).

The tier-specific patterns of distribution of LBs aggregates were preserved across the whole rostrocaudal extent of the SNc in all biological specimens (Figure 6B), although slight changes in the ratio between pigmented neurons with and without LB burden were found in the female animal with eight months in-life (M307f8), the one showing a more severe nigrostriatal damage.

## DISCUSSION

Parallel trajectories between the observed tier-specific patterns of intracellular pigmentation levels and LBs burden were described here. Neurons located in SNcV are the ones showing the highest levels of pigmentation together with the greatest abundance of intracellular inclusions, whilst CB+ SNcD neurons accounted for the opposite (e.g., low pigmentation levels and a minimal number of LBs). It is worth noting that the intraparenchymal deliveries of AAV1-*hTyr* were performed just dorsal to the SNc, and therefore SNcD neurons should have been the ones more heavily neuromelanized, however this was not the case. Indeed, the highest pigmentation levels were constantly found in SNcV neurons. Although the expression of the *hTyr* gene was under the control of a ubiquitous promoter (CMV), only catecholaminergic neurons had the chance to accumulate NMel following tyrosinase enhancement, therefore here the specific transduction of dopaminergic neurons was driven by the encoded transgene, instead of promoter-related^4^. The direct link between pigmentation levels and intracellular inclusions was consistent across all experimental subjects, even considering that less than a half of the total number of TH+ neurons in the SNc became neuromelanized upon the delivery of the AAV1-*hTyr* vector. Moreover, the conducted quantification of LBs is inherently underestimated, resulting from any given number of TH+ pigmented neurons that had already degenerated at the time of the necropsy. Although the underlying mechanisms for the well-known tier-specific dopaminergic cell vulnerability within the context of PD remain to be elucidated in full detail, data reported here provided evidence for a close association between high pigmentation levels, presence of LBs and lack of CB as three synergistic players sustaining a combination of relevant risk factors triggering dopaminergic cellular degeneration.

To date, up to 20 different clusters of dopaminergic cell subpopulations have been reported within the ventral mesencephalon, comprising the SNc, the ventral tegmental area, and the retrorubral area^22^, well beyond the basic dichotomy between Aldh1a1+ and CB+ neurons. Different cellular clusters have been identified on the basis of their specific neurochemical profiles, expression levels, projection patterns, and vulnerability vs. resilience to degeneration. In brief, at the level of the SNcV, diverse clusters of TH+ neurons expressing Aldh1a1 with or without Girk2, Sox6, and Anxa1 co-expression have been reported, whereas SNcD contains neurons positive for CB and vGlut2, the latter more predominant in laterodorsal SNc territories^16,21–24^. Moreover, identified neurochemical profiles also exhibit different projection patterns, where nigrostriatal projections mainly arise from Aldh1a1+ neurons in the SNcV, whilst CB+ neurons located in SNcD preferentially innervate extrastriatal territories such as the internal and external divisions of the globus pallidus^12,16^. Despite this vast heterogeneity, when compared to more resilient SNcD CB+ dopaminergic cells, SNcV neurons expressing Aldh1a1 have often been taken as the most vulnerable ones^25^, in particular those co-expressing angiotensin II receptor type 1^26^, matching the degeneration patterns observed in human PD^27,28^.

Over the past few decades, a prominent role of synuclein aggregation process has been broadly accepted in the field of PD research and even as a key concept in clinical practice to explain a clinico-pathological correlate of disease progression^29,30^. However, it is worth recognizing that pathology is not pathophysiology, and LBs may be considered as a natural reaction of dopaminergic neurons against a number of insults rather than the causative triggers of neurodegeneration^31–33^. Indeed, the fact that LBs were found within pigmented neurons staying alive at the time of necropsy in human postmortem PD samples (the same applies to pigmented rodent and NHP models of PD)^4,5^, raises some concerns about the extent to which these intracytoplasmic aggregates of phosphorylated α-Syn can be regarded as truly detrimental for dopaminergic neurons^34,35^.

When searching for other possible mechanisms and factors sustaining dopaminergic cell neurodegeneration, NMel may be regarded as a good candidate to be appointed as “the butler”, and indeed, the data provided here, together with available cumulative evidence, are pointing in this direction, and from different complementary perspectives and datasets. Although initial hypotheses linking NMel and dopamine cell vulnerability had been postulated a long time ago^36–38^, this association was later neglected since rodents, as the most often used experimental biological specimens, lack NMel pigmentation in dopaminergic neurons, and for unknown reasons^6^. Moreover, a link between age-related NMel pigmentation as the inducer of synucleinopathy and subsequent vulnerability of dopaminergic neurons has been postulated^34^. Furthermore, conducted analyses in postmortem PD brain samples revealed that A9 neurons are predisposed to aggregate α-Syn around pigment-associated lipids under conditions of oxidation and iron loading, reactions known to precipitate α-Syn^35^. Furthermore, tonic production of dopamine depends on oxidative mechanisms^39–41^, and indeed NMel is the end by-product of the non-enzymatic auto-oxidation of dopamine^42–45^. Blaming NMel for PD-related degeneration of A9 neurons is also supported by the proven bidirectional clinical association between the incidences of melanoma (a cancer of skin pigmented cells) and PD^46–52^. Likewise, α-Syn has been detected in cultured melanoma cells and in malignant melanocytes^53^. Another piece of evidence was gathered from IPSC-derived dopaminergic cells taken from PD patients harboring mutations in the *PARK7* gene, leading to loss-of-function of DJ-1, a protective protein against oxidative stress^61^. When compared to IPSC-derived dopaminergic neurons from patients with different PD-related mutations (LRKK2 and GBA1), only those neurons derived from *PARK7* mutations exhibited spontaneous pigmentation at p50^55^. Finally, different gene therapy-based modalities intended to reduce NMel levels^3,56^ have resulted in dopaminergic neuroprotection together with a reduction of LBs burden.

In conclusion, the data presented here confirm that the pigmented model of PD in macaques expresses pathological features typically observed at the level of the SNcV in postmortem brain samples of human PD and therefore, could be conveniently used to further understand the beginning of nigrostriatal neurodegeneration and to test potential disease-modifying therapeutics.

## METHODS

### Study design

This study took advantage of NHP brain samples already available from a previous study^4^ where an adeno-associated viral vector serotype 1 coding for the hTyr gene was delivered into the left substantia nigra in four NHPs (*Macaca fascicularis*). The conducted study was intended to assess a potential correlation between tier-specific vulnerability of dopaminergic neurons, the distribution patterns of LBs within the SNc, and pigmentation levels in NHPs. Experiments were carried out in four experimental subjects (two males and two females), euthanized four and eight-months post-injection of the viral vector (M308f4 and M319m4 vs. M307f8 and M309m8, respectively). Animals numbered M307f8 and M308f4 were females, whereas animals M309m8 and M310m4 were both males. LB-like intracytoplasmic inclusions were mapped throughout the whole extent of the SNc, where dorsal and ventral tiers were identified by the expression of calbindin and aldehyde dehydrogenase type 1a1 proteins (CB and Aldh1a1, respectively). Moreover, intracellular NMel levels were measured at the single-cell level in identified neurons located within SNcD and SNcV tiers, in an attempt to correlate pigmentation levels and LB pathology.

### Experimental subjects

A total of four adult juvenile naïve macaques (*Macaca fascicularis;* 36-40 months old; two males and two females; body weight 2.3-4.5 Kg) were used in this study. Animal handling was conducted in accordance with the European Council Directive 2010/63/UE as well as in keeping with the Spanish legislation (RD53/2013). Animal experimentation was approved by the Ethical Committee for Animal Testing of the University of Navarra (ref: CEEA095/21) as well as by the Department of Animal Welfare of the Government of Navarra (ref: 222E/2021).

### Viral vectors

Recombinant AAV vector serotype 2/1 containing the human tyrosinase cDNA under the control of a CMV promoter (AAV-hTyr) and the corresponding control empty vector (AAV-null) were produced at the Viral Vector Core Production Unit of the Autonomous University of Barcelona (www.viralvector.eu). Details on viral vector synthesis, plasmids, and sequences, can be found elsewhere (Chocarro et al., 2023).

### Ventriculography-assisted stereotactic surgery for AAV deliveries

Surgical anesthesia was induced by intramuscular injection of ketamine and midazolam (5 mg/Kg each). Local anesthesia was implemented just before surgery with a 10% solution of lidocaine. Analgesia was achieved with a single intramuscular injection of flunixin meglumine (Finadyne®, 5 mg/Kg), delivered at the end of the surgery and repeated 24 and 48 h post-surgery. A similar schedule was conducted for antibiotic coverage (ampicillin, 0.5 ml/day). Once the experimental subjects showed a complete recovery, they were returned to the animal vivarium and housed in groups.

Stereotactic coordinates for the SNc were calculated from the atlas of Lanciego and Vázquez^57^, and target selection was assisted by intrasurgical ventriculographies (sagittal projections) confirming the right placement of the injection device (a Hamilton® syringe) before AAV deliveries into either the left and right SNc. Injections were performed in pulses of 1 μl/min for a total volume of 10 μl each in two sites of the SNc, each deposit spaced 1 mm in the rostrocaudal axis in an attempt to transduce the whole extent of the SNc. Once viral deliveries were completed, the needle was left in place for an additional 10 min before withdrawal to minimize reflux of the AAV suspensions through the injection tract. Coordinates were 5.5 mm caudal to the anterior commissure (ac), 5 mm ventral to the bicommissural plane, and 4 mm lateral to the midline for the rostral deposits of AAV-*hTyr* (left SNc) and AAV-null (right SNc), whereas the caudal injections were placed 8.5 mm caudal to the ac; 5.5 mm ventral to the ac-pc plane, and 4 mm lateral to the mid-sagittal vein sinus.

#### Necropsy and tissue processing

Animals numbered M308f4 and M310m4 were euthanized after four months in-life, whereas a follow-up time of eight months was applied to specimens identified as M307f8 and M309m8. Terminal anesthesia was first induced with an intramuscular injection of ketamine (10 mg/Kg), followed by an overdose of sodium pentobarbital (200 mg/Kg). Animals were perfused with a saline Ringer solution (500 ml) followed by 3,000 ml of a fixative made of 4% paraformaldehyde and 0.1% glutaraldehyde in 0.125 M phosphate buffer, pH 7.4. Perfusion was continued with 1,000 ml of a cryoprotectant buffered solution containing 10% glycerin and 1% dimethylsulphoxide (DMSO). Once the perfusion was completed, the brain was removed from the skull and stored for 48 h in a cryoprotectant solution made of 20% glycerin and 2% DMSO in 0.125 M phosphate buffer at neutral pH. Next, ten series of frozen coronal brain adjacent sections (40 μm-thick) were obtained on a sliding microtome, each series being used for different histochemical and immunohistochemical (immunoperoxidase or immunofluorescent) stains.

#### Quantification of pigmented neurons in the SNc

To calculate a ratio between pigmented and non-pigmented neurons, every 10^th^ section was counterstained with neutral red, digitized at a magnification of 20x (Aperio SC2 scanner, Leica, Wetzlar, Germany), and uploaded to the Aiforia® cloud (www.aiforia.com) for image analysis. Within this platform, a dedicated bi-layered algorithm was prepared and validated (error of 1.65%), disclosing between pigmented neurons (NR+ NMel+) and non-pigmented neurons (NR+ NMel-). Two additional series of sections were processed for the single immunoperoxidase detection of aldehyde dehydrogenase type 1a1 (Aldh1a1) and calbindin (CB), scanned at a magnification of 20x, and analyzed with specific algorithms quantifying the number of pigmented neurons in the ventral and dorsal tiers of the SNc (Aldh1a1+ NMel+ and CB+ NMel+, respectively). Furthermore, intracellular NMel levels were quantified with Fiji ImageJ (NIH, USA) at the single-cell level in five equally-spaced sections covering the whole rostrocaudal extent of the SNc and stained for either Aldh1a1 or CB according to available protocol^58^. Tier-specific analyses of intracellular pigmentation levels were conducted in 631 pigmented neurons in animal M307f8, 670 neurons in M308f4, 891 neurons in M309m8, and 997 neurons in M310m8.

### Lewy body-like burden, quantification, and tier-specific distribution

Every 10^th^ section was processed for the multiple immunofluorescent detection of markers characteristic of Lewy body-like inclusions (LBs). Accordingly, stains disclosing phosphorylated alpha-synuclein (pSer-129) and P62 were combined under the confocal laser-scanning microscope with the immunofluorescent detection of tyrosine hydroxylase (TH; disclosing dopaminergic neurons) and brightfield visualization of NMel. Furthermore, two more series of sections were stained for Aldh1a1, P62, and TH and for CB, P62, and TH, to further evaluate the precise tier-specific distribution of LB burden.

### Statistical analysis

Statistical analyses were performed with GraphPad Prism version 10.1.2. for Windows and Stata 16 (StataCorp LLC, College Station, TX, USA). Relevant tests are listed in the corresponding figure legends. Species with *p* < 0.05 were considered statistically significant.

## DECLARATIONS

### Availability of data and materials

Further information and requests for resources and reagents should be directed to and will be fulfilled by the corresponding author, Jose L. Lanciego. The data, protocols, and key lab materials used and generated in this study are listed in a Key Resource Table alongside their persistent identifiers at (https://doi.org/10.5281/zenodo.18936255). No code was generated for this study; all data cleaning, preprocessing, analysis, and visualization were performed using Fiji ImageJ, GraphPad Prism 10.1.2, Stata16, and Aiforia.

### Competing interests

The authors report no competing of interest

## Acknowledgements

This research was funded by MICIU/AIE/10.13039/501100011033 (Grant No. PID2023-127802OB-I00) and FEDER, UE.

## Authors’ contributions

Conceptualization, AJR, JB, JAO, JLL; methodology, JC, GA, ELR, MMI, CC, AL-V; data analysis, JC, AJR, MMI, CC, AL-V; visualization, JC, AJR, GA, ELR, MMI, CC, JB; funding acquisition, JLL; resources, AJR, GA, JLL; project administration, AJR, JLL; supervision, writing – original draft, JC, AL-V, AJR, JLL; writing – reviewing & editing, final draft, JC, JB, JAO, JLL.

## List of abbreviations

AAV: adeno-associated virus
Aldh1a1: aldehyde dehydrogenase type 1a1
α-Syn: alpha-synuclein
Anxa1: Annexin A1
CB: calbindin
CMV: cytomegalovirus
DJ-1: protein encoded by the PARK7 gene
DMSO: dimethylsulphoxide
GBA1: beta-glucocerebrosidase gene 1
Girk2: G-protein-activated inward rectifying potassium channel type 2
hTyr: Human tyrosinase gene
IPSCs: induced pluripotent stem cells
LBs: Lewy body-like
LRRK2: leucine-rich repeat kinase 2
NHPs: non-human primates
NMel: neuromelanin
OD: optical density
PD: Parkinson’s disease
SNc: substantia nigra pars compacta
SNcD: substantia nigra pars compacta, dorsal tier
SNcV: substantia nigra pars compacta, ventral tier
TH: tyrosine hydroxylase
vGlut2: vesicular glutamate transporter type 2

## Notes

https://doi.org/10.5281/zenodo.18936255

### Competing Interest Statement

The authors have declared no competing interest.

